# Tomato immune receptor Ve1 recognizes surface-exposed co-localized N- and C-termini of *Verticillium dahliae* effector Ave1

**DOI:** 10.1101/103473

**Authors:** Yin Song, Zhao Zhang, Jordi C. Boshoven, Hanna Rovenich, Michael F. Seidl, Jernej Jakše, Karunakaran Maruthachalam, Chun-Ming Liu, Krishna V. Subbarao, Branka Javornik, Bart P.H.J. Thomma

**Affiliations:** Laboratory of Phytopathology, Wageningen University, Droevendaalsesteeg 1, 6708 PB Wageningen, The Netherlands.; Key Laboratory of Plant Molecular Physiology, Institute of Botany, the Chinese Academy of Sciences, Beijing 100093, China.; Center for Plant Biotechnology and Breeding, University of Ljubljana, Jamnikarjeva 101, 1000 Ljubljana, Slovenia.; Department of Plant Pathology, University of California, Davis, CA 95616, USA; Present address: Department of Ornamental Horticulture and Landscape Architecture, China Agricultural University, Beijing 100193, China.

## Abstract

Effectors are secreted by plant pathogens to facilitate infection, often through deregulation of host immune responses. During host colonization, race 1 strains of the soil-borne vascular wilt fungus *Verticillium dahliae* secrete the effector protein Ave1 that triggers immunity in tomato genotypes that encode the Ve1 immune receptor. Homologs of *V. dahliae* Ave1 (VdAve1) are found in plants and in few plant pathogenic microbes, and are differentially recognized by Ve1. However, how VdAve1 is recognized by Ve1 remained unknown. Interestingly, C-terminally affinity-tagged versions of VdAve1 failed to activate Ve1-mediated immunity, suggesting that exposure of the C-terminus of VdAve1 is required for Ve1-mediated recognition. This was confirmed by subsequent analysis of C-terminal deletion mutants, and by domain swap experiments. Although required, only the C-terminus of VdAve1 is not sufficient to activate Ve1-mediated immunity. Intriguingly, a three-dimensional structural model of VdAve1 revealed that the N- and C-termini co-localize on a surface-exposed patch of the VdAve1 protein. Indeed, subsequent analyses of N-terminal deletion mutants confirmed that also the N-terminus of VdAve1 is required to activate Ve1-mediated immunity. Thus, we conclude that a surface-exposed patch of the VdAve1 protein that is composed by co-localized N- and C-termini is recognized by the tomato immune receptor Ve1.

## INTRODUCTION

Plants are constantly engaged in battles against diverse groups of microbes within their environment. However, only few of these actually become pathogens to cause disease, as plants have developed innate immunity to protect themselves against microbial attack (Dodds & Rathjen, 2010; Thomma, BPHJ et al., 2011; Cook et al., 2015). In its simplest form, plant immunity against pathogen attack is governed by immune receptors that sense pathogen-derived(induced) ligands to activate defense. Originally, the interaction between plant immune receptors and pathogen ligands was described in the “gene-for-gene” model, stating that the products of plant resistance (*R*) genes induce race-specific resistance upon recognition of the products of corresponding pathogen avirulence (*Avr*) genes (Flor, 1971). Decades later, an updated view of plant innate immunity has been introduced as the “zigzag” model, by incorporating pathogen-secreted effector molecules that suppress host immune responses, but that may subsequently be recognized by newly evolved immune receptors, in turn (Jones & Dangl, 2006). In this model, the first line of defense is governed by plasma membrane-localized pattern recognition receptors (PRRs) that detect conserved microbe-associated molecular patterns (MAMPs) to activate MAMP-triggered immunity (MTI). In subsequent layers of defense, effectors are recognized by the corresponding resistance proteins (R proteins), resulting in effector-triggered immunity (ETI). Although initially portrayed as separate layers, numerous studies on various plant-microbe interactions have revealed that the delineation between MTI and ETI is not strict, but rather a continuum (Thomma et al., 2011; Cook et al., 2015). Moreover, the fact that MAMPs are defined from the perspective of the host, whereas effectors are defined from the perspective of the invader, creates a conceptual conflict and has recently inspired the formulation of the Invasion Model, in which host receptors (termed invasion pattern receptors; IPRs) detect either externally encoded or modified-self ligands (termed invasion patterns; IPs) that betray invasion (Cook et al., 2015). In this model, any molecule can serve as an IP that is detected by an IPR, but the probability of a particular ligand-receptor complex to evolve within the framework of host immunity increases with increasing ligand probability to retain function, conservation across organisms, importance in establishment of symbiosis, and accessibility (Cook et al., 2015).

*Verticillium dahliae* is a xylem invading fungal pathogen that causes Verticillium wilt diseases in a wide range of plant species worldwide (Fradin & Thomma, 2006). *V. dahliae* persists in the soil and enters plants through their roots. Once inside the root, the fungus grows intercellularly and invades the xylem vessels, where it sporulates to spread through the vascular system. Typical symptoms of *V. dahliae* infection include stunting, wilting, chlorosis, and necrosis (Fradin & Thomma, 2006). In tomato (*Solanum lycopersicum*), a single dominant locus that confers *Verticillium* resistance has been identified as the *Ve* locus, which controls *Verticillium* isolates that are assigned to race 1 (Schaible et al., 1951). The *Ve* locus comprises two closely linked and inversely oriented genes, *Ve1* and *Ve2*, that both encode extracellular leucine-rich repeat (eLRR) receptor-like proteins (RLPs) (Kawchuk et al., 2001; Wang et al., 2010). Of these, only *Ve1* was found to confer resistance against race 1 isolates of *Verticillium* in tomato (Fradin et al., 2009). Intriguingly, interfamily transfer of *Ve1* from tomato to *Arabidopsis thaliana* has resulted in race-specific *Verticillium* resistance in the latter species (Fradin et al., 2011; 2014; Zhang et al., 2014), implying that the underlying immune signalling cascade across plant taxonomy is evolutionarily conserved (Fradin et al., 2011; Thomma et al., 2011). Moreover, homologs of tomato Ve1 that act as immune receptors against race 1 strains of *V. dahliae* have been characterized in other plant species including tobacco, potato, wild eggplant and hop, suggesting an ancient origin of the tomato immune receptor Ve1 (Song et al., 2016).

Through comparative population genomics, the *V. dahliae* effector protein that is recognized by the tomato immune receptor Ve1 was identified as Ave1 (for <u>A</u>virulence on <u>Ve1</u> tomato) (de Jonge et al., 2012). Although the intrinsic function of *V. dahliae* Ave1 (VdAve1) remains elusive, it is clear that VdAve1 contributes to fungal virulence on susceptible plant genotypes (de Jonge et al., 2012). Interestingly, homologs of VdAve1 were identified from a number of fungal pathogens, including the tomato pathogen *Fusarium oxysporum* f. sp. *lycopersici* (FoAve1), the sugar beet pathogen *Cercospora beticola* (CbAve1) and the Brassicaceae pathogen *Colletotrichum higginsianum* (ChAve1) (de Jonge et al., 2012). Strikingly, however, most VdAve1 homologs were found in plants, with the most closely related homologs derived from tomato (*S. lycopersicum*; SlPNP) and grape (*Vitis vinifera*; VvPNP) (de Jonge et al., 2012). Finally, a more distantly related homolog was identified in the plant pathogenic bacterium *Xanthomonas axonopodis* pv. *citri* (XacPNP) (de Jonge et al., 2012). Co-expression of *VdAve1*, *SlPNP*, *FoAve1*, and *CbAve1* with tomato *Ve1* in tobacco triggers a hypersensitive response (HR), whereas co-expression of *ChAve1* with tomato *Ve1* does not lead to an HR (de Jonge et al., 2012). Moreover, Ve1 was found to mediate resistance towards *F. oxysporum* in tomato, demonstrating involvement of the tomato immune receptor Ve1 in resistance against multiple fungal pathogens (de Jonge et al., 2012).

Many eLRR-containing cell-surface immune receptors recognize peptide sequences as epitopes of their pathogen ligands. For example, flg22 is the 22-amino-acid peptide derived from bacterial flagellin that is perceived by the receptor-like kinase (RLK)-type immune receptor FLS2 (Zipfel et al., 2004), while the *Arabidopsis* RLK-type EFR immune receptor was shown to recognize elf18, an 18-amino-acid peptide derived from bacterial EF-Tu (Zipfel et al., 2006).

Similarly, a highly conserved 22-amino-acid sequence derived from the bacterial cold shock protein, named csp22, is perceived by the tomato RLK-type immune receptor CORE (Felix & Boller, 2003; Wang et al., 2016), while a surface-exposed pentapeptide TKLGE of the 22 kDa ethylene-inducing xylanase (EIX) from the biocontrol fungus *Trichoderma viride* determines recognition by the RLP-type receptor LeEIX2 in tomato (Rotblat et al., 2002; Ron & Avni, 2004). Furthermore, the tyrosine-sulfated 21-amino-acid peptide RaxX21-sY of *Xanthomonas oryzae* pv. *oryzae* RaxX is sufficient to activate XA21-mediated immunity in rice (Pruitt et al., 2015). Finally, a conserved 20-amino-acid fragment present in most Nep1-like proteins (NLPs) (nlp20) is sufficient to activate RLP23-mediated immunity in *Arabidopsis* (Böhm et al., 2014; Oome et al., 2014; Albert et al., 2015). Here, we aimed to identify a minimal motif in the *V. dahliae* effector protein VdAve1 that is necessary and sufficient for recognition by the tomato immune receptor Ve1. Our approach was based on epitope prediction, guided by the alignment of differentially recognized VdAve1 homologs, followed by functional analyses of a combination of serial deletion assays, domain swaps, synthetic peptides, three-dimensional structural prediction and chimeric proteins.

## RESULTS

### Sequence conservation among Ave1 homologs

Previously, we reported the cloning of *Ave1* from *V. dahliae* and described the absence of allelic variation among 85 *Ave1* alleles derived from *V. dahliae* and *V. albo-atrum* (presently *V. alfalfae*; Inderbitzin et al., 2011) (de Jonge et al., 2012). *Ave1* alleles were not identified in any of the *V. dahliae* and *V. albo-atrum* race 2 strains analysed, nor in strains that are not pathogenic on tomato, nor in *V. longisporum* or *V. tricorpus* (de Jonge et al., 2012). To further explore *Ave1* diversity, we assessed presence in a collection of 129 *Verticillium* strains isolated from various host plants and different geographical locations, resulting in the identification of 22 novel *Ave1* alleles (Table S1). No allelic variation was found among the newly identified *Ave1* alleles amplified from *V. dahliae* as well as from *V. alfalfae* and *V. nonalfalfae*, two species that have recently been recognized as novel species distinct from *V. albo-atrum* (Inderbitzin et al., 2011). However, a *VdAve1* homolog was identified in four isolates of *V. nubilum* (*VnAve1*), a species that is known as a saprophyte and opportunistic pathogen (Isaac, 1953). While the predicted VnAve1 protein sequence displays 13 amino acid polymorphisms when compared with VdAve1 (Figure 1A), the four *VnAve1* alleles were found to be identical to each other.

**Figure 1.**
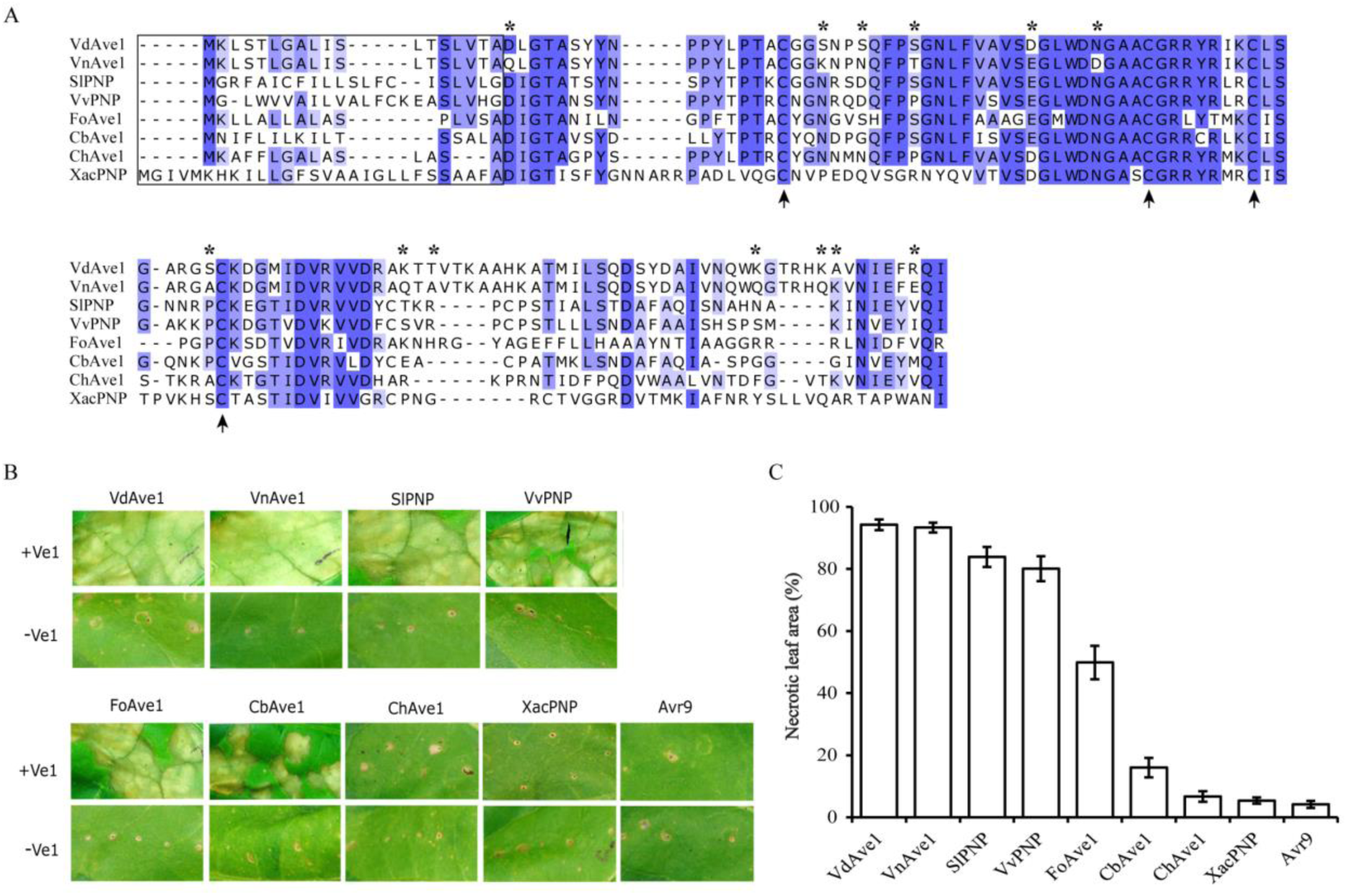
Distinct necrosis induced by *VdAve1* homologs through co-expression with tomato *Ve1* in *Nicotiana tabacum*. (A) Amino acid sequence alignment of VdAve1 homologs from *Verticillium dahliae* (VdAve1), *V. nubilum* (VnAve1), *Solanum lycopersicum* (SlPNP), *Vitis vinifera* (VvPNP), *Fusarium oxysporum* f. sp. *lycopersici* (FoAve1), *Cercospora beticola* (CbAve1), *Colletotrichum higginsianum* (ChAve1), and *Xanthomonas axonopodis* pv. citri (XacPNP). Blue shade background indicates identical amino acids while the color intensity represents the frequency. Asterisks indicate the 13 amino acid polymophisms between VdAve1 and VnAve1. The positions of four conserved cysteine residues are indicated with arrows in the bottom. The N-terminal amino acids in the frame denote the predicted signal peptides (SP) of the VdAve1 homologs. (B) Coexpression of *Ve1* and *Ave1* homologs VdAve1, VnAve1, SlPNP, VvPNP, FoAve1, CbAve1, ChAve1, and XacPNP in *N. tabacum*. Expression of the sequence-unrelated effector *Avr9* from the tomato leaf mold fungus *Cladosporium fulvum* in combination with *Ve1* or absence of *Ve1* are shown as negative controls. Pictures were taken at five days post infiltration (dpi). (C) Quantification of necrosis resulting from recognition of VdAve1 homologs by Ve1 at 5 dpi (n > 20). Bars represent the average percentage of necrotic leaf area of infiltration zones with standard deviations.

Alignment of the amino acid sequences of VdAve1 and its homologs from plants (SlPNP and VvPNP) and plant pathogens (VnAve1, FoAve1, CbAve1, ChAve1 and XacPNP) revealed blocks of highly conserved amino acids that are alternated with more variable regions (Figure 1A). Based on prediction by SignalP 4.0 (Petersen et al., 2011), all VdAve1 homologs contain a predicted N-terminal signal peptide that directs secretion into the extracellular space (Figure 1A; D-cutoff score >0.6). Moreover, the four cysteine residues that are present in VdAve1 are conserved among all homologs (Figure 1A), and *in silico* analysis using DISULFIND (Ceroni et al., 2006) suggests the formation of disulphide bridges between Cys^35^ and Cys^63^, as well as between Cys^71^ and Cys^79^. From the alignment it is apparent that XacPNP is the most divergent, while all other VdAve1 homologs are relatively similar (Figure 1A).

### Comparison of the necrosis-inducing activity of VdAve1 homologs

It was previously demonstrated that tomato Ve1 recognizes not only VdAve1, but also SlPNP, FoAve1 and CbAve1 (de Jonge et al., 2012). We now also tested the necrosis-inducing capacity of VnAve1, VvPNP and XacPNP that were isolated from *V. nubilum, V. vinifera* and *X. axonopodis*, respectively. Co-expression of the sequence-unrelated effector *Avr9* from the tomato leaf mold fungus *Cladosporium fulvum* (van Kan et al., 1991) with *Ve1* served as a negative control. Whereas expression of *VnAve1* or *VvPNP* together with *Ve1* in *Nicotiana tabacum* resulted in strong HR, co-expression of *XacPNP* or *Avr9* with *Ve1* triggered little to no necrosis in addition to the small wounds that were caused by the infiltration procedure (Figure 1B). To compare the necrosis induced by the various VdAve1 homologs, they were co-expressed with *Ve1* in *N. tabacum* and HR development was measured by quantification of the leaf area that developed necrosis at five days post infiltration (dpi) (Zhang et al., 2013). Importantly, none of the VdAve1 homologs induced necrosis in the absence of Ve1 (Figure 1B). Whereas agroinfiltration of either *VdAve1* or *VnAve1* with *Ve1* resulted in complete necrosis of the infiltrated leaf area, agroinfiltration of *FoAve1* with *Ve1* resulted in large necrotic spots in the infiltrated leaf area, although no complete collapse of the infiltrated area was observed (Figure 1B; C). Upon agroinfiltration of *CbAve1* with *Ve1*, spreading of smaller and larger necrotic spots was observed in all infiltrated areas, but the infiltrated leaf area did not turn completely necrotic. For *ChAve1*, *XacPNP* and *Avr9*, necrosis did not extend beyond the wounded infiltration sites (Figure 1B; C). Upon agroinfiltration of the tomato and grape *VdAve1* homologs, *SlPNP* and *VvPNP*, most of the infiltrated leaf area developed necrosis (Figure 1B; C), occasionally affecting the complete infiltrated leaf sector. To confirm that variable levels of HR induced by VdAve1 homologs are not due to instability of the VdAve1 homologs *in planta*, GFP-tagged VdAve1 homologs were detected by immunoblotting. Similar to GFP-tagged VdAve1 protein or GFP-tagged VnAve1, all other GFP-tagged VdAve1 homologs accumulated to clearly detectable protein levels *in planta* (Figure S1).

### The C-terminus of VdAve1 is required, but not sufficient, for recognition by Ve1

In order to perform further functional analyses, a construct encoding C-terminally GFP-tagged VdAve1 (VdAve1_c GFP) was generated. However, C-terminal fusion of a GFP tag to VdAve1 resulted in loss of recognition by Ve1 (Figure 2A; C). Considering that the GFP tag (~27 kDa) is relatively large, we engineered C-terminally tagged VdAve1 fusions with smaller protein tags (<12 kDa; VdAve1_c 3xHA, VdAve1_c 6xHis, VdAve1_c 4xMyc, VdAve1_c 10xMyc and VdAve1_c FLAG; Figure 2A). Despite their smaller size, all additionally tested C-terminal tags abolished, or significantly reduced, HR development on *Ve1*-expressing tobacco leaves (Figure 2A; C). Importantly, the C-terminally GFP-tagged VdAve1 fusion was clearly detected by immunoblotting (Figure S2A), suggesting that accessibility of the VdAve1 C-terminus is important in VdAve1 recognition by Ve1.

**Figure 2.**
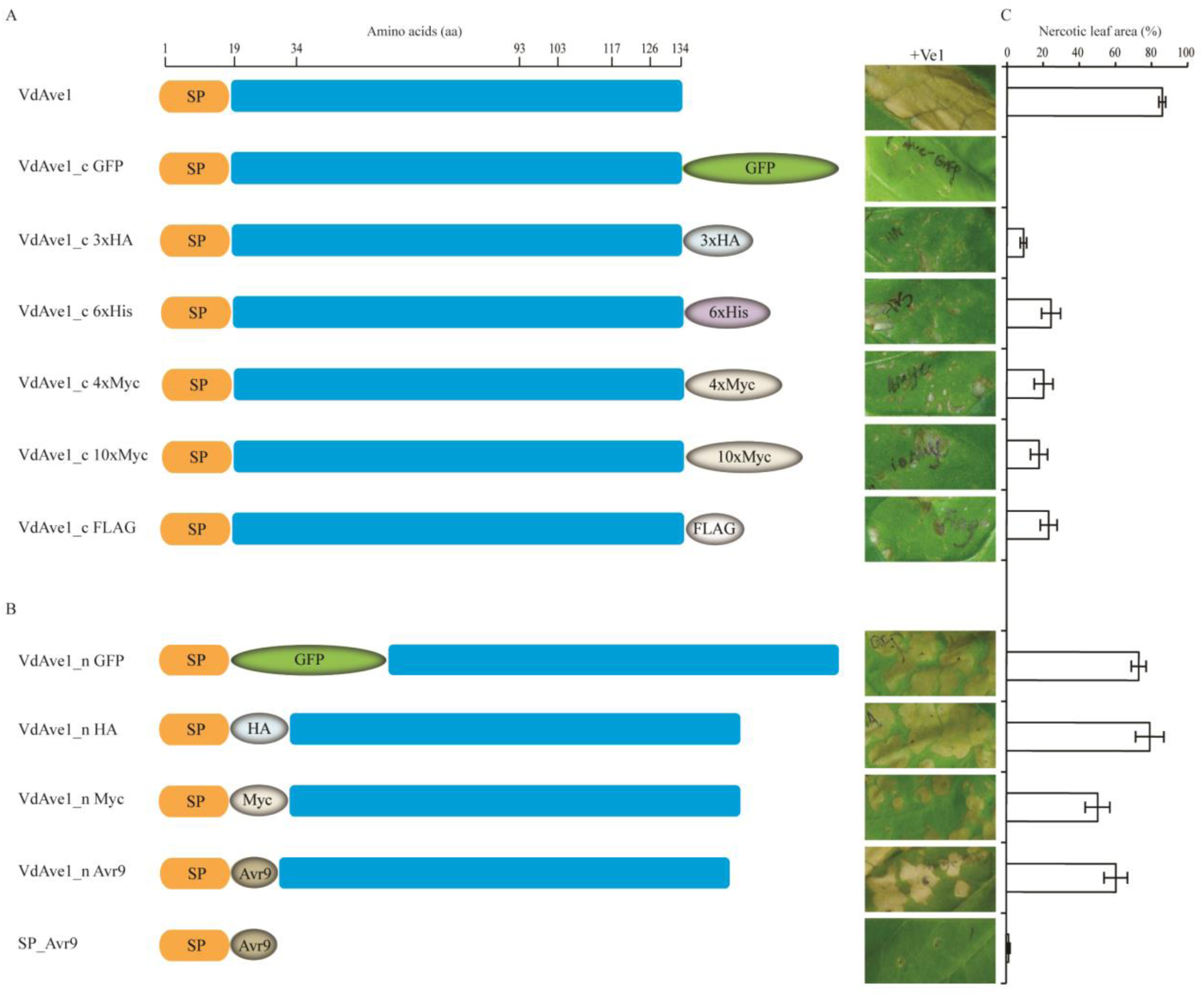
Effect of C- or N- terminally tagged VdAve1 versions on recognition of VdAve1 by Ve1 in tobacco. (A) Constructs encoding full-length VdAve1 (VdAve1) and C-terminally tagged VdAve1 versions (VdAve1_c GFP, VdAve1_c 3xHA, VdAve1_c 6xHis, VdAve1_c 4xMyc, VdAve1_c 10xMyc, VdAve1_c FLAG) were assayed for necrosis-inducing capability. The constructs were co-expressed with *Ve1* in tobacco respectively, and the occurrence of necrosis was monitored at 5 dpi. The feature includes a predicated signal peptide (SP) involved in effector secretion. (B) Constructs encoding N-terminally fused VdAve1 versions (VdAve1_n GFP, VdAve1_n HA, VdAve1_n Myc and VdAve1_n Avr9) were assayed for necrosis-inducing capability. The constructs were co-expressed with *Ve1* in tobacco respectively, and the necrosis occurrence was monitored at 5 dpi. A construct encoding mature Avr9 fused with VdAve1 signal peptide (SP_Avr9) co-expressed with *Ve1* is used as a negative control. (C) Quantification of necrosis resulting from recognition of tagged VdAve1 proteins by Ve1 at 5 dpi. The graph shows the average percentage of necrotic leaf area of infiltration zones at 5 dpi (n > 5). Data are presented as mean with standard deviations.

To further investigate the role of the C-terminus in recognition of VdAve1 by Ve1, a number of C-terminal truncation mutants were generated. Deletion of 42 amino acids from the C-terminus (Lys^93^ to Ile^134^; VdAve1^Δ93-134^) resulted in loss of VdAve1 recognition by Ve1 (Figure 3A; B). Subsequent analysis of step-wise smaller truncations revealed that a C-terminal deletion of nine amino acids (VdAve1^Δ126-134^) resulted in loss of Ve1-mediated recognition (Figure 3A; B), even though the presence of GFP-tagged VdAve1^Δ126-134^ protein *in planta* was verified by immunoblotting (Figure S2A). We subsequently performed complementation experiments in *V. dahliae* to confirm the importance of the C-terminus for activation of Ve1-mediated immunity. To this end, we expressed *VdAve1^Δ126-134^* driven by the native *VdAve1* promoter in a *VdAve1* deletion mutant (*V. dahliae* Δ*VdAve1*; de Jonge et al., 2012) and inoculated *Ve1* tomato plants with the complemented strains. Plants that were inoculated with three independent *V. dahliae* strains expressing *VdAve1*^*Δ126-134*^ (VdAve1^Δ126-134^ #1, VdAve1^Δ126-134^ #2 and VdAve1^Δ126-134^ #3) showed a similar disease phenotype as *Ve1* plants inoculated with the *V. dahliae* JR2 Δ*VdAve1* strain, whereas plants inoculated with wild-type *V. dahliae* strain JR2 and the *VdAve1* complementation strain resembled mock-inoculated plants (Figure 3C; D). Collectively, these results demonstrate that the C-terminal nine amino acids of VdAve1 are required to activate Ve1-mediated immunity.

**Figure 3.**
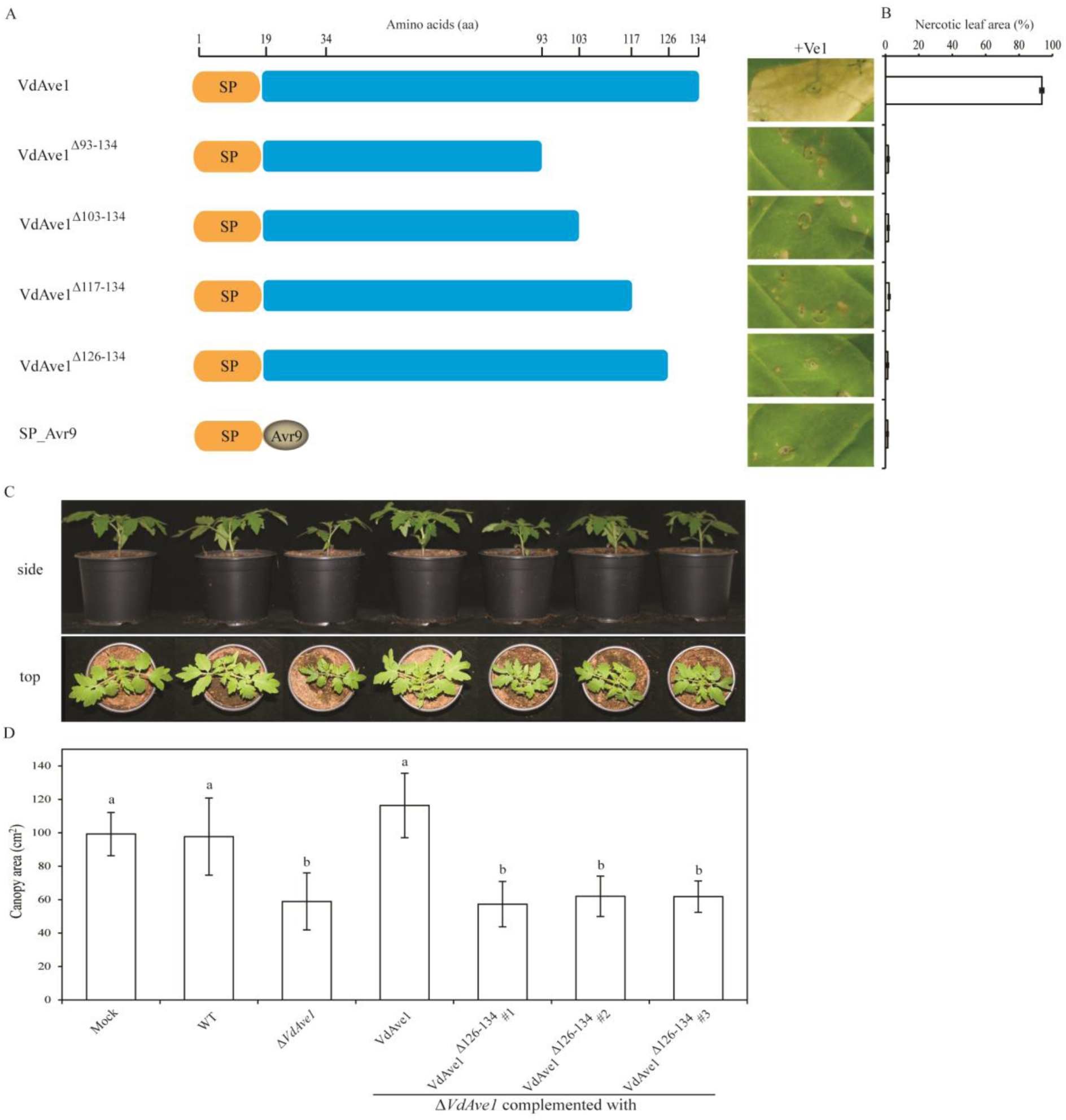
The C-terminal nine amino acids of VdAve1 are required to establish Ve1-mediated immunity. (A) VdAve1 C-terminal truncations result in loss of recognition by Ve1. Constructs encoding four VdAve1 truncations that lack the C-terminal 42 (VdAve1^Δ93-134^), 32 (VdAve1^Δ103-134^), 18 (VdAve1^Δ117-134^) and 9 (VdAve1Δ^126-134^) amino acids of VdAve1 co-infiltration with *Ve1* were assayed respectively, and the occurrence of necrosis was recorded at 5 dpi. Constructs VdAve1 and SP_Avr9 were used as a positive and negative control, respectively. The feature includes a predicated signal peptide (SP) involved in effector secretion. (B) Quantification of necrosis resulting from recognition of VdAve1 C-terminal truncations by Ve1 at 5 dpi. The graph shows the average percentage of necrotic leaf area of infiltration zones at 5 dpi (n > 5). Data are presented as mean with standard deviations. (C) Complementation assays in *Verticillium dahliae* show that the VdAve1 C-terminal nine amino acids are required to activate Ve1-mediated immunity in tomato. Three independent *V. dahliae VdAve1* deletion (*ΔVdAve1*) strains expressing a construct encoding VdAve1 lacking the C-terminal nine amino acids (VdAve1Δ^126-134^ #1, VdAve1Δ^126-134^ #2 and VdAve1Δ^126-134^ #3) escape recognition by *Ve1* tomato compared with *V. dahliae* wild-type (WT) and genetic complementation strains (VdAve1) evidenced by stunted *Ve1* plants at 14 days post *Verticillium* inoculation. (D) Average canopy area of 8 *Ve1* tomato plants inoculated with different *V. dahliae* strains or mock-inoculation. Different letters indicate significant differences (*P* < 0.05). The data shown are representative of three independent experiments.

Since the C-terminal nine amino acids appear to be essential for VdAve1 recognition, and the bacterial homolog XacPNP that is significantly divergent in this region (Figure 1A) is not recognized by Ve1 (Figure 1B; C), C-terminal nine-amino-acid swaps between VdAve1 and XacPNP were performed. An expression construct encoding a chimeric VdAve1 protein was engineered in which the C-terminal nine amino acids of VdAve1 were replaced by those of XacPNP (Vd_XacPNP^c 9AA^; Figure 4A). As expected, similar to XacPNP, co-expression of *Vd_XacPNP*^*c 9AA*^ with *Ve1* in tobacco failed to induce an HR, as only minimal necrotic spots were observed (Figure 4A; B). Conversely, a construct encoding another chimeric VdAve1 protein was generated in which the last nine amino acids of XacPNP were replaced by those of VdAve1 (Xac_VdAve1^c 9AA^; Figure 4A). Co-expression of *Xac_VdAve1*^*c 9AA*^ with *Ve1* in tobacco induced a relatively strong HR when compared with XacPNP or Vd_XacPNP^c 9AA^-induced HR, although full necrosis was not observed in the infiltrated leaf area (Figure 4A; B). Immunoblotting confirmed that the GFP-tagged Vd_XacPNP^c 9AA^ and Xac_VdAve1^c 9AA^ proteins were present *in planta* (Figure S2B). These results confirm that the C-terminal nine amino acids of VdAve1 are critical for recognition by Ve1, and furthermore suggest that these are sufficient for recognition.

**Figure 4.**
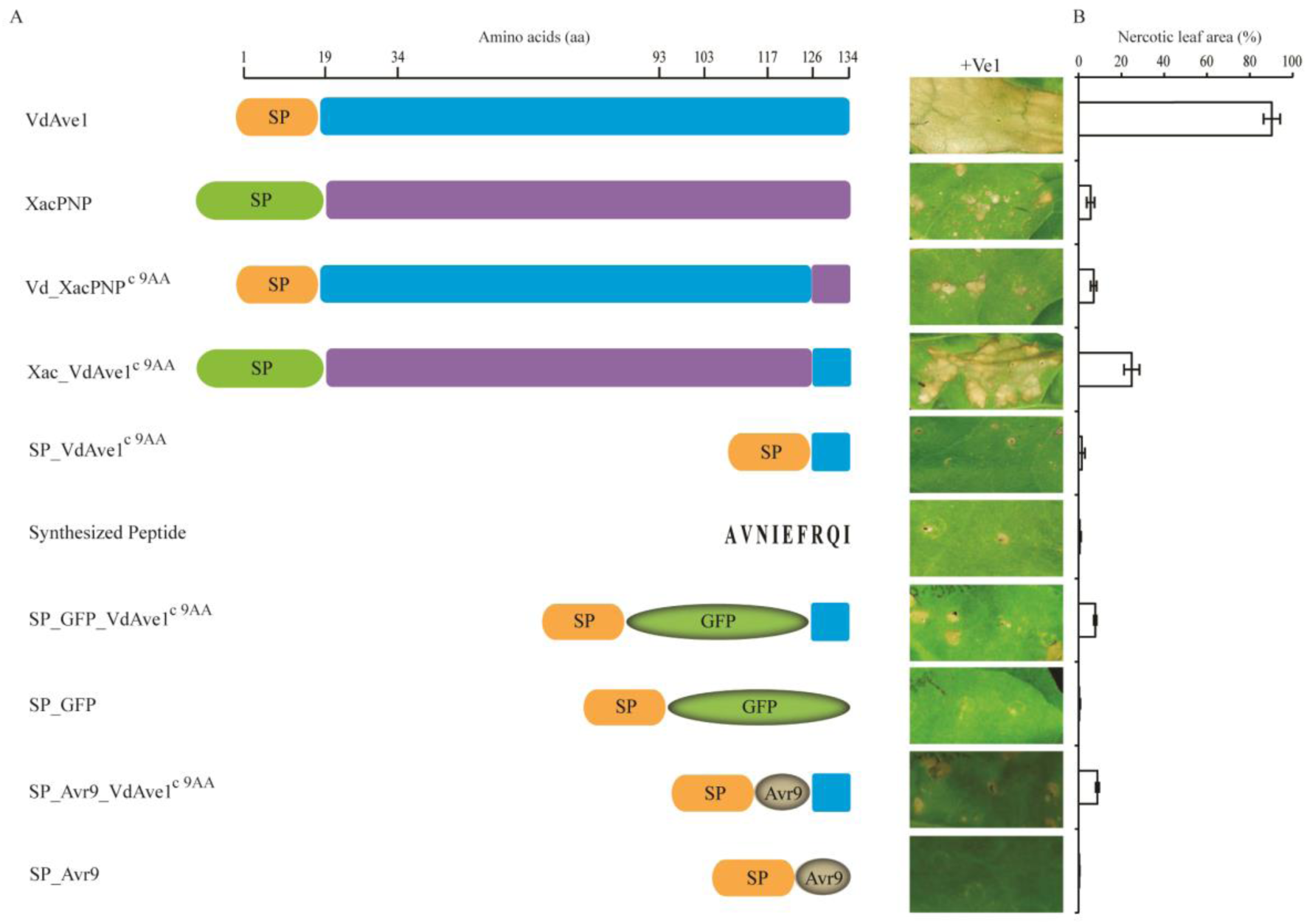
(A) The C-terminal nine amino acids of VdAve1 are critical but not sufficient for Ve1-mediated recognition in tobacco. Constructs encoding full-length VdAve1 and XacPNP with their own signal peptides (SP) and two chimeras in which their C-terminal nine amino acids were swapped (Vd_XacPNP^c 9AA^ and Xac_VdAve1^c 9AA^) were assayed. In chimera Vd_XacPNP^c 9AA^ the C-terminal nine amino acids of VdAve1 were replaced by those of XacPNP, while in chimera Xac_VdAve1^c 9AA^ the C-terminal nine amino acids of XacPNP were replaced by those of XacPNP. A construct encodes the C-terminal nine amino acids of VdAve1 fused to the VdAve1 signal peptide (SP_VdAve1^c 9AA^). A construct encodes a GFP that is C-terminally fused to the C-terminal nine amino acids of VdAve1, and N-terminally fused to the VdAve1 signal peptide to establish extracellular secretion (SP_GFP_VdAve1^c 9AA^), while a construct encoding GFP fused with the VdAve1 signal peptide (SP_GFP) and construct SP_Avr9 were used as negative controls. Furthermore, a chemically synthetized peptide encompassing VdAve1 C-terminal nine amino acids peptide (AVNIEFRQI) was used. A construct VdAve1 was used as a positive control. The feature includes a predicated signal peptide (SP) involved in effector secretion. All the constructs were co-expressed with *Ve1* in tobacco respectively, and the occurrence of necrosis was monitored at 5 dpi. (B) Quantification of necrosis resulting from recognition of VdAve1 chimeras by Ve1 at 5 dpi. The graph shows the average percentage of necrotic leaf area of infiltration zones at 5 dpi (n > 10). Data are presented as mean with standard deviations.

To determine whether the C-terminal nine amino acids are indeed sufficient to trigger Ve1-mediated recognition, we generated a construct encoding the C-terminal nine amino acids of VdAve1 fused to the VdAve1 signal peptide (SP_VdAve1^c 9AA^; Figure 4A). This construct was co-expressed with *Ve1* in tobacco, but no necrosis was observed in the infiltrated leaf (Figure 4A; B). Additionally, infiltration of a chemically synthetized peptide encompassing the C-terminal nine amino acids of VdAve1 (peptide sequence: AVNIEFRQI) did not induce an HR in *Ve1*-expressing tobacco up to a concentration of 1 mg/mL (Figure 4A; B). However, possibly the nine amino acid peptide is not stable in the apoplast. In an attempt to overcome such complications, we generated two constructs in which the coding sequence of GFP or Avr9 was N-terminally fused to the VdAve1 signal peptide and C-terminally fused to the C-terminal nine amino acids of VdAve1 (SP_GFP_VdAve1^c 9AA^ and SP_Avr9_VdAve1^c 9AA^; Figure 4A). As negative controls, two constructs in which the coding sequence of GFP or Avr9 was N-terminally fused to the VdAve1 signal peptide without the C-terminal nine amino acids of VdAve1 (SP_GFP and SP_Avr9; Figure 4A) were generated. All constructs were co-expressed with *Ve1* in tobacco and HR development was monitored at 5 dpi. However, only slight necrosis was observed following infiltration of *Ve1* with *SP_GFP_VdAve1*^*c 9AA*^ or *SP_Avr9_VdAve1*^*c 9AA*^ in the infiltrated sector (Figure 4A; B), despite detection of SP_GFP_VdAve1^c 9AA^ protein *in planta* by immunoblotting (Figure S2A). These data reveal that the C-terminal nine amino acids of VdAve1 are not sufficient to activate *Ve1*-mediated immunity.

To test whether longer stretches of the C-terminus of VdAve1 can be used to trigger Ve1-mediated recognition, we generated a construct encoding the C-terminal 18 amino acids of VdAve1 fused to the VdAve1 signal peptide (SP_VdAve1^c 18AA^; Figure 5A). This construct was co-expressed with *Ve1* in tobacco, but only slightly increased necrosis was observed in the infiltrated sector (Figure 5A; B). We further generated two constructs in which the coding sequence of GFP or Avr9 was N-terminally fused to the VdAve1 signal peptide and C-terminally fused to the C-terminal 18 amino acids of VdAve1 (SP_GFP_VdAve1^c 18AA^ and SP_Avr9_VdAve1^c 18AA^; Figure 5A). Co-infiltrations of *Ve1* with *SP_GFP* or *SP_Avr9* were used as negative controls. Co-expression of *Ve1* with *SP_GFP_VdAve1*^*c 18AA*^ or *SP_Avr9_VdAve1*^*c 18AA*^ in tobacco induced similar necrosis formation as SP_VdAve1^c 18AA^ protein at 5 dpi (Figure 5A; B). Immunoblotting confirmed that the chimeric SP_GFP_VdAve1^c 18AA^ protein was present *in planta* (Figure S3). Subsequent analyses showed that also the C-terminal 32 amino acids and the C-terminal 42 amino acids of VdAve1 induce signs of a weak HR at 5 dpi (Figure 5A; B), although their presence was verified by immunoblotting (Figure S3). Taken together, these results reveal that the C-terminus of VdAve1 is required, but not sufficient, to activate Ve1-mediated recognition.

**Figure 5.**
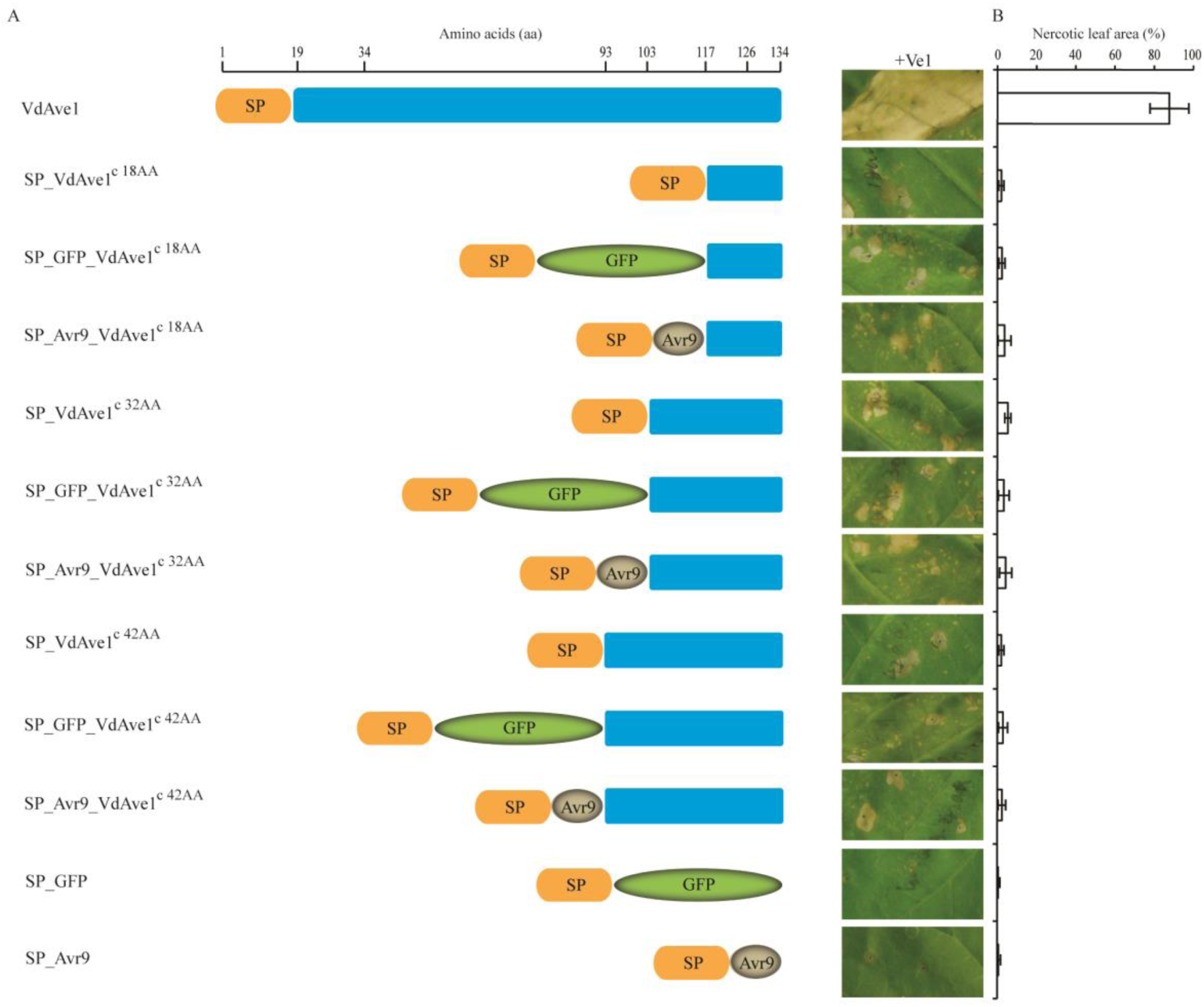
VdAve1 C-terminus is not sufficient for VdAve1 recognition by Ve1. (A) Occurrence of the necrosis in tobacco upon co-expression of VdAve1 C-terminal chimeras with *Ve1*. Three constructs encoding the VdAve1 signal peptide (SP) fused to the VdAve1 C-terminal 18-amino-acid (SP_VdAve1^c 18AA^), or the GFP and VdAve1 C-terminal 18-amino-acid (SP_GFP_ VdAve1^c18AA^), or the mature Avr9 and VdAve1 C-terminal 18-amino-acid (SP_Avr9_ VdAve1^c 18AA^) were generated. Furthermore, constructs encoding the extended VdAve1 C-termini (SP_VdAve1^c32AA^ and SP_VdAve1^c 42AA^), or N-terminally GFP-tagged VdAve1 C-terminal extensions (SP_GFP_VdAve1^c 32AA^ and SP_GFP_VdAve1^c 42AA^), or N-terminally Avr9-fused VdAve1 C-terminal extensions (SP_Avr9_VdAve1^c 32AA^ and SP_Avr9_VdAve1^c 42AA^) were assayed too. Constructs VdAve1, SP_GFP and SP_Avr9 were used as controls. The feature includes a predicated signal peptide (SP) involved in effector secretion. The constructs were co-expressed with *Ve1* in tobacco respectively, and the necrosis occurrence was recorded at 5 dpi. (B) Quantification of necrosis resulting from recognition of VdAve1 C-terminal chimeras by Ve1 at 5 dpi. The graph shows the average percentage of necrotic leaf area of infiltration zones at 5 dpi (n > 10). Data are presented as mean with standard deviations.

### Structural modelling reveals co-localization of VdAve1 C- and N-termini on a surface-exposed patch

In an attempt to gain more insight in VdAve1 recognition by Ve1, a three-dimensional structural model of VdAve1 was generated. Structural comparison with the protein databank (RCSB PDB) (Rose et al., 2013) revealed the maize protein EXPB1 (PDB ID: 2HCZ) (Yennawar et al., 2006) as a potential structural analogue (TM-Score 0.84) of VdAve1. The VdAve1 structural model shows that the C-terminal nine amino acids (Figure 6; shown in red) are exposed at the surface of the mature VdAve1 protein. The model also predicts that the N-terminus (Figure 6; shown in orange) congregates with the C-terminus on the exposed surface of the mature VdAve1 protein (Figure 6), which suggests that Ve1 may recognize a patch of the VdAve1 protein that includes both the N- and C-termini.

**Figure 6.**
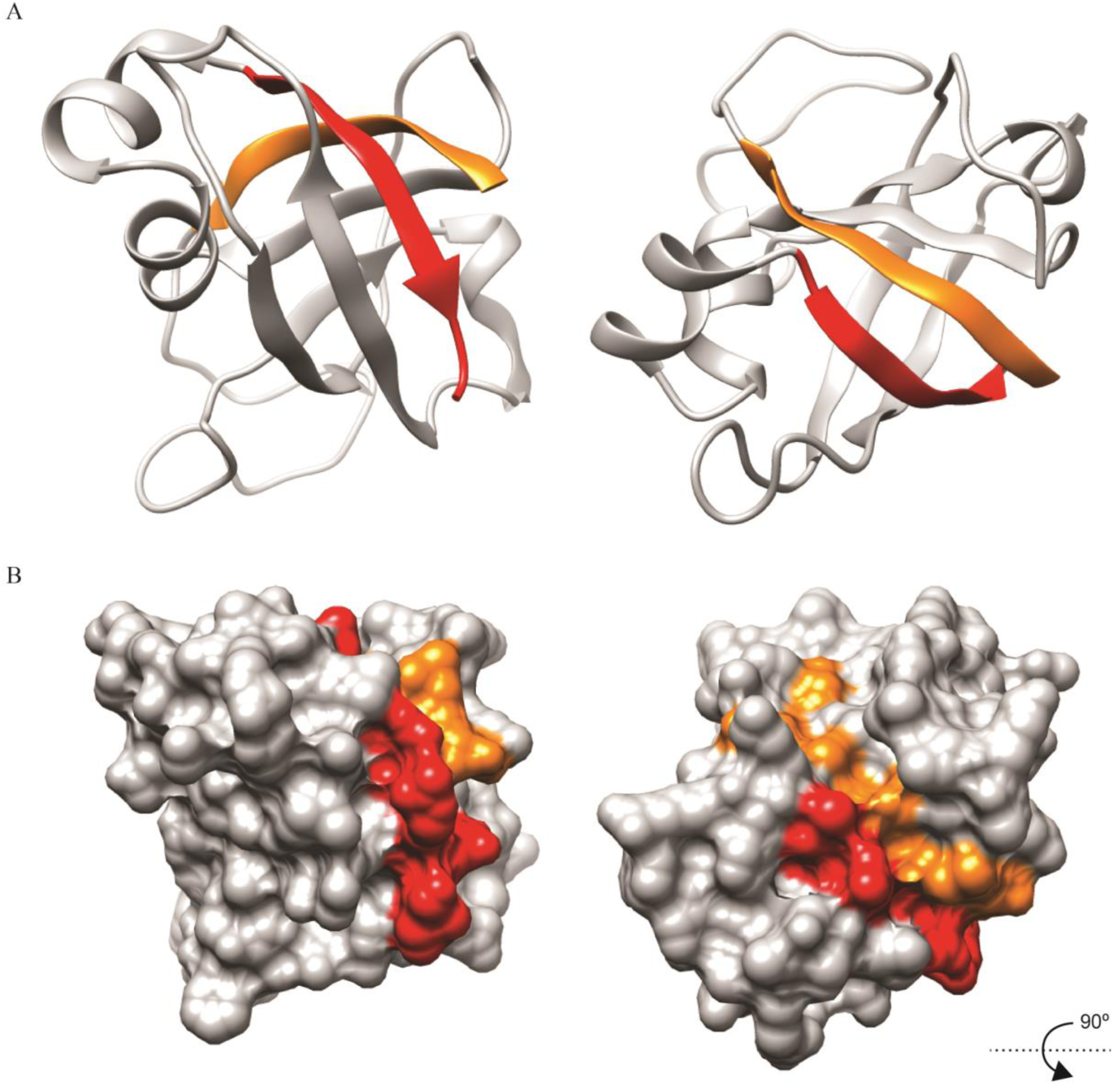
Three-dimensional structural model of the VdAve1 protein. The VdAve1 structure was predicted using I-TASSER (Zhang, 2008). The inferred VdAve1 structure is of high quality indicated by a confidence score (C-Score) of 1.22, and displayed as a ribbon (A), and surface (B) model from the side (left) and the top (right). The C-terminal nine amino acids sequence of VdAve1 is indicated in red, while the N-terminal eight amino acids sequence is indicated in orange.

### The N-terminus of VdAve1 is required, but not sufficient for Ve1-mediated recognition

We previously observed that affinity tags that were C-terminally fused to VdAve1 compromised recognition by Ve1, resulting in the finding that the C-terminus of VdAve1 is required for recognition. To test whether we could implement the N-terminus in recognition by Ve1 too, we fused various tags (GFP, HA, Myc and Avr9) to the N-terminus of mature VdAve1. Indeed, we similarly observed compromised Ve1-mediated recognition upon use of the Myc (VdAve1_n Myc) or Avr9 (VdAve1_n Avr9) tag (Figure 2B; C), even though N-terminal tagging of VdAve1 with GFP (VdAve1_n GFP) or HA (VdAve1_n HA) did not clearly affect recognition by Ve1 (Figure 2B; C). Immunoblotting confirmed that the N-terminally GFP-tagged VdAve1 protein was stably produced *in planta* (Figure S2A). These data suggest that, besides the VdAve1 C-terminus, also the N-terminus of VdAve1 is involved in recognition by Ve1.

To further investigate the importance of the N-terminus of VdAve1 in Ve1-mediated recognition, recognition of VdAve1 upon deletion of the N-terminal 16 amino acids (from Asp^19^ to Ala^34^; VdAve1^Δ19-34^) by Ve1 was tested in tobacco. Indeed, deletion of the 16 N-terminal amino acids of the mature VdAve1 abolished recognition by Ve1 (Figure 7A; B). Furthermore, we engineered two constructs in which the coding sequence of GFP or Avr9 was N-terminally fused to the VdAve1 signal peptide and C-terminally fused to the mature VdAve1 lacking the N-terminal 16-amino-acid of mature VdAve1 (SP_GFP_VdAve1^Δ19-34^ and SP_Avr9_ VdAve1^Δ19-34^; Figure 7A). Co-expression of *Ve1* with *SP_GFP_ VdAve1*^*Δ19-34*^ or *SP_Avr9_ VdAve1*^*Δ19-34*^ in tobacco did not result in Ve1-mediated recognition (Figure 7A; B), although production of SP_GFP_VdAve1^Δ19-34^ was confirmed by immunoblotting (Figure S4). To further confirm the involvement of the VdAve1 N-terminus in Ve1-mediated recognition, we introduced the coding sequence *VdAve1*^*Δ19-34*^ under the control of the native *VdAve1* promoter into *V. dahliae* JR2 *ΔVdAve1* (de Jonge et al., 2012) and performed a disease assay on *Ve1* tomato plants This assay showed that, similar to *V. dahliae* JR2 *ΔVdAve1*, also complementation strains expressing *VdAve1*^*Δ19-34*^ caused clear Verticillium wilt symptoms on *Ve1* tomato plants, while wild-type *V. dahliae* strain JR2 and the complementation strain expressing *VdAve1* caused no disease symptoms on tomato plants expressing *Ve1* (Figure 7C; D). These results reveal that the N-terminal 16 amino acids of mature VdAve1 are required to trigger Ve1-mediated immunity.

**Figure 7.**
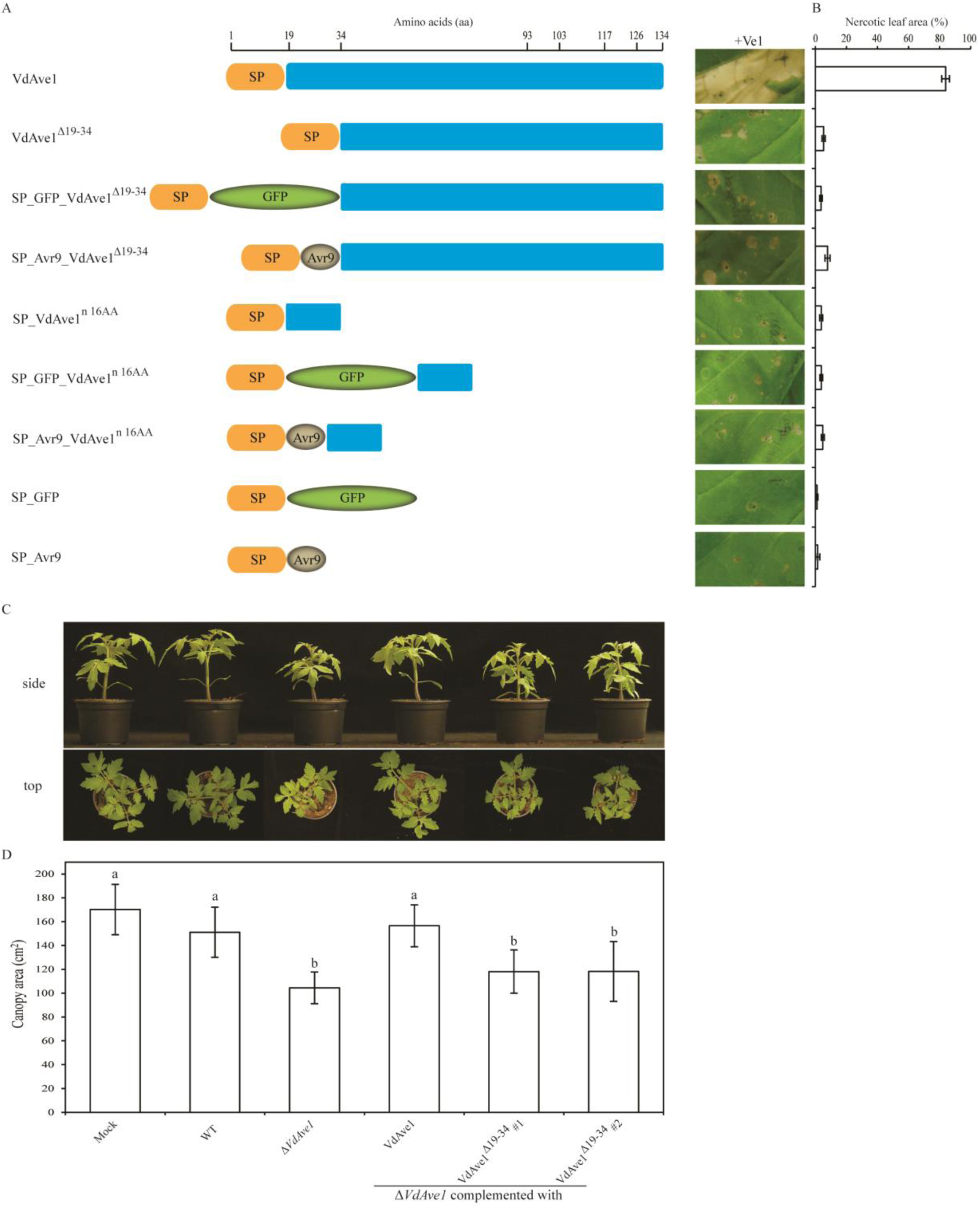
(A) The N-terminal sixteen amino acids of mature VdAve1 are required but not sufficient for Ve1-mediated recognition in tobacco. Occurrence of the necrosis in *N. tabacum* upon co-expression of N-terminal chimeras with *Ve1*. Constructs encoding a VdAve1 truncation that lacks the N-terminal 16-amino-acid of mature VdAve1 (VdAve1^Δ19-34^), and the corresponding N-terminal GFP or Avr9 fusion (SP_GFP_VdAve1^Δ19-34^ and SP_Avr9_VdAve1^Δ19-34^) were used. Furthermore, constructs encoding the VdAve1 signal peptide (SP) fused to the N-terminal 16-amino-acid of mature VdAve1 (SP_VdAve1^n 16AA^), or the GFP and the N-terminal 16-amino-acid of mature VdAve1 (SP_GFP_VdAve1^n 16AA^), or the mature Avr9 and VdAve1 C-terminal 16-amino-acid (SP_GFP_VdAve1^n 16AA^) were assayed. Constructs VdAve1, SP_GFP and SP_Avr9 were used as controls. The constructs include a predicted signal peptide (SP) to direct effector secretion. The constructs were co-agroinfiltrated with *Ve1* in tobacco respectively, and the necrosis occurrence was monitored at 5 dpi. (B) Quantification of necrosis resulting from recognition of VdAve1 N-terminal chimeras by Ve1 at 5 dpi. The graph shows the average percentage of necrotic leaf area of infiltration zones at 5 dpi (n > 10). Data are presented as mean with standard deviations. (C) Complementation assays in *V. dahliae* show that the N-terminal sixteen amino acids of mature VdAve1 protein are required to activate Ve1-mediated immunity in tomato. Two independent *V. dahliae VdAve1* deletion (Δ*VdAve1*) strains expressing a construct encoding VdAve1 lacking the N-terminal sixteen amino acids of mature VdAve1 (VdAve1 Δ^19-34^ #1 and VdAve1 Δ^19-34^ #2) escape recognition by *Ve1* tomato compared with *V. dahliae* wild-type (WT) and genetic complementation strains (VdAve1) evidenced by stunted *Ve1* plants at 14 days post *Verticillium* inoculation. (D) Average canopy area of 8 *Ve1* tomatoplants inoculated with different *V. dahliae* strains or mock-inoculation Different letters indicate significant differences (*P* < 0.05). The data shown are representative of three independent assays.

To confirm that the N-terminal 16 amino acids are not sufficient to trigger Ve1-mediated recognition, we engineered a construct encoding the N-terminal 16-amino-acid of mature VdAve1 fused to the VdAve1 signal peptide (SP_VdAve1^n 16AA^; Figure 7A). Furthermore, two constructs were designed in which the coding sequence of GFP or Avr9 was N-terminally fused to the VdAve1 signal peptide and C-terminally tagged to the N-terminal 16-amino-acid of mature VdAve1 (SP_GFP_VdAve1^n 16AA^ and SP_Avr9_VdAve1^n 16AA^; Figure 7A). Co-expression of *Ve1* with none of these constructs in tobacco induced an HR (Figure 7A; B), although immunoblotting showed that the SP_GFP_VdAve1^n 16AA^ was present *in planta* (Figure S4). Collectively, these results demonstrate that the N-terminal 16 amino acids of mature VdAve1 are required, but not sufficient, for establishment of Ve1-mediated immunity.

## DISCUSSION

We have previously shown that the *V. dahliae* effector VdAve1 is recognized by the tomato immune receptor Ve1 (de Jonge et al., 2012), and we have identified several homologs of VdAve1 that are differentially recognized by Ve1 (Figure 1B; C). In this study, we demonstrated that a surface-exposed patch of the VdAve1 protein that is composed of co-localized N- and C-termini of *V. dahliae* effector VdAve1 is recognized by tomato immune receptor Ve1. Our analyses reveal that the C-terminus as well as the N-terminus individually are required, but not sufficient, to activate Ve1-mediated immunity.

Plant cell surface-localized PRRs are often activated upon recognition of short peptide sequences on the surface of their ligands, such as flg22 or flgII-28 that are derived from flagellin (Felix et al., 1999; Cai et al., 2011; Clarke et al., 2013), the pentapeptide TKLGE derived from EIX (Rotblat et al., 2002), the csp22 peptide derived from the bacterial cold shock proteins (Felix & Boller, 2003), elf18 or EFa50 derived from EF-Tu (Kunze et al., 2004; Furukawa et al., 2014), and the nlp20 peptide found in most NLPs (Böhm et al., 2014; Oome et al., 2014). In our study, we attempted to identify such motif in VdAve1. Clearly, our results suggest that the co-localization of the two termini of the primary amino acid sequence, rather than a contiguous stretch of amino acids composes the recognition motif. Our efforts to identify an artificial minimal peptide that can be recognized by tomato Ve1 by generating chimeric peptides consisting of the N-terminal sixteen amino acids in combination with various C-terminal peptides, and fused to various tags, were fruitless (data not shown). This suggests that also the folding of the recognized sequence patch, and the spatial orientation of the two protein termini, is important for the activation of Ve1-meidated immunity.

Although several minimal motifs have been identified in ligands of cell surface receptors, similar examples for intracellular NLR (nucleotide-binding domain leucine-rich repeat) immune receptors have not been reported. Moreover, it has suggested that simultaneous recognition of multiple epitopes within a single effector is required for NLR activation. For example, distinct regions of the *Pseudomonas syringae* effector AvrRps4 are required for the activation of the intracellular NLR receptors PRS4/RRS1-mediated immunity (Sohn et al., 2009; 2012). Similarly, multiple contact points are likely required for recognition of the flax-rust effectors AvrL567 and AvrM by the corresponding NLR receptors in flax (Wang et al., 2007; Ve et al., 2013). Likewise, multiple residues at separate locations on the surface of the *Hyaloperonospora arabidopsidis* effector ATR1 are required for recognition by the *Arabidopsis* NLR receptor RPP1 (Chou et al., 2011; Goritschnig et al., 2016). In contrast to these findings, our data suggest that, although physically separated on the primary amino acid chain, recognition converges at a single surface-exposed patch of the VdAve1 protein.

Thus far, we have failed to show a direct physical interaction between VdAve1 and Ve1. Functional analysis of the tomato immune receptor Ve1 through domain swaps with its non-functional homolog Ve2, and subsequent alanine scanning mutagenesis on the solvent exposed β-strand/β-turn residues across the eLRR domain previously identified several regions of the Ve1 protein that are required for functionality (Fradin et al., 2014; Zhang et al., 2014). In these studies, Ve1 functionality was restricted to three consecutive eLRR regions, namely eLRR1-eLRR8, eLRR20-eLRR23 and eLRR32-eLRR37, of which two regions eLRR1-eLRR8 and eLRR20-eLRR23 were proposed to contribute to ligand binding, while eLRR32-eLRR37 was proposed to function in immune signalling activation (Zhang et al., 2014). Realistically, final confirmation of this model cannot be obtained through domain swaps, domain deletions, gene shuffling analyses and site-directed mutagenesis within the immune receptor or the recognized ligand, but will ultimately have to follow from structural analysis of receptor-ligand interactions, for instance through crystallography.

## MATERIALS AND METHODS

### Plant materials and plant growth conditions

Tobacco (*Nicotiana tabacum* cv. Petite Havana SR1) and *35S::Ve1* tomato (*Solanum lycopersicum* cv. MoneyMaker background; Fradin et al., 2009) plants were grown in the greenhouse (Unifarm, Wageningen, the Netherlands) at 21°C/19°C during 16/8 hours day/night periods, respectively, with 70% relative humidity and 100 W/m^−2^ supplemental light when the light intensity dropped below 150 W/m^−2^. After agroinfiltration, tobacco plants were grown in the climate room at 22°C/19°C during 16-h/8-h day/night periods, respectively, with 70% relative humidity.

### Generation of binary expression vectors

Construction of all binary expression vectors is described in Methods S1.

### Transient expression assays

Overnight cultures of *A. tumefaciens* strain GV3101 containing expression constructs were harvested at OD600 of 0.8 to 1 by centrifugation and resuspended to a final OD of 2 in infiltration medium as described previously (Zhang et al., 2013). *A. tumefaciens* cultures containing constructs to express Ve1 and VdAve1, or VdAve1 homologs, or VdAve1 chimeras were mixed in a 1:1 ratio and infiltrated into leafs of five- to six-week-old tobacco plants. At five days post infiltration (dpi), infiltrated leaves were photographed, and necrosis was quantified by using ImageJ to measure the area of necrosis as percentage of the total infiltrated leaf area (Song et al., 2016).

### Protein extracts and immunoblotting

For immunological detection of GFP-tagged VdAve1 homologs and VdAve1 chimeras, *A. tumefaciens* carrying the corresponding expression constructs was infiltrated into mature tobacco leaves as described previously (Zhang et al., 2013). The co-immunopurifications and immunoblotting were performed as described previously (Zhang et al., 2014).

### Generation of complementation *V. dahliae* strains

To generate *VdAve1*, *VdAve1*^*Δ126-134*^ and *VdAve1*^*Δ19-34*^ complementation constructs, DNA fragments containing the *Pac*I and *Not*I restriction sites were amplified by PCR from the corresponding plasmids VdAve1, VdAve1^Δ126-134^ and VdAve1^Δ19-34^ using primers listed in Table S2, and cloned into the vector pFBT005 which *ToxA* promoter was replaced by the native *VdAve1* promoter (~1.6 kb) and contains a nourseothricin cassette, respectively. All the constructs were confirmed by DNA sequencing (Eurofins Genomics, Ebersberg, Germany), and subsequently transformed into *A. tumefaciens* strain AGL1 by electroporation. *A. tumefaciens*-mediated transformation of *V. dahliae* strain JR2 *ΔVdAve1* (de Jonge et al., 2012) was performed as previously described (Santhanam, 2012). *V. dahliae* transformants were selected on potato dextrose agar (PDA; Oxoid, Basingstoke, UK) plates containing 50 µg/mL nourseothricin sulphate (Sigma-Aldrich Chemie BV, Zwijndrecht, the Netherlands). After five to seven days at room temperature, individual transformants were transferred to fresh PDA plates and incubated for 7 to 10 days. Genomic DNA was extracted from individual transformants and PCR was performed to test presence of the inserted nourseothricin cassette and presence of the inserted chimeric *VdAve1* fragment.

### Disease assays

*V. dahliae* was grown on PDA plates at 22 °C, and conidia were collected from 7- to 10- day-old *V. dahliae* cultures on PDA plates and washed with tap water. Disease assays on tomato plants were performed as previously described (Fradin et al., 2009). Briefly, 10-day-old tomato plants were uprooted, the roots were rinsed in water, dipped for 5 min in a suspension of 10^6^ conidiospores/mL water while the roots of mock plants were dipped in tap water without conidiospores, and transplanted to fresh commercial potting soil (Horticoop, Bleiswijk, the Netherlands). Disease symptoms were scored up to 14 days post *Verticillium* inoculation, inoculated plants were photographed. The canopy area of 8 plants was measured with ImageJ software and a One-Way ANOVA was performed with IBM SPSS statistics software.

### Generation of the structural model of VdAve1

The *V. dahliae* VdAve1 structure was predicted using I-TASSER v4.3 (Zhang, 2008) and rendered using UCSF Chimera v1.10.1 (Pettersen et al., 2004). Structural predictions with C-Scores > −1.5 are generally considered to have a correct fold (C-Scores are typically in the range of [-5,2]) (Roy et al., 2010). The structural analog in the protein data bank (RCSB PDB) (Rose et al., 2013) was identified using the TM-align program which is part of the I-TASSER package. Analogous structures with TM-Scores > 0.5 are considered to have a similar fold (TM-Scores in the range [0,1]) (Roy et al., 2010).

## ACKNOWLEDGEMENTS

Y.S. acknowledges a PhD fellowship from the China Scholarship Council (CSC). B.P.H.J.T. and M.F.S are supported by a Vici and a Veni grant, respectively, of the Research Council for Earth and Life sciences (ALW) of the Netherlands Organization for Scientific Research (NWO). We thank Bert Essenstam and Pauline Sanderson at Unifarm for excellent plant care, and Sebastjan Radišek at the Institute of Hop Research and Brewing, Slovenia, for providing *Verticillium* strains.

## AUTHOR CONTRIBUTIONS

Y.S, Z.Z., C.-M. and B.P.H.J.T. planned and designed the research. Y.S, Z.Z., J.C.B., H.R., M.F.S., J.J. and K.M. performed experiments, Y.S, Z.Z., J.C.B., H.R., M.F.S., J.J., K.V.S., B.J. and B.P.H.J.T. analysed data. Y.S, Z.Z., and B.P.H.J.T. wrote the manuscript. All authors read and approved the final manuscript.

